# Effects of complex whole-body movements on EEG activity: a scoping review

**DOI:** 10.1101/2025.03.13.642812

**Authors:** Gabriel Müller, Atef Salem, Wolfgang I. Schöllhorn

## Abstract

The influence of movements on brain activity has been part of research for decades. Recent advancements in electroencephalography (EEG) coupled with a shift in focus towards the effects of complex whole-body movements provided additional inspirations in this area. Besides the metabolic load, the amount of information to be processed in parallel provides rough indications of its influence on central nervous activity. Accordingly, this scoping review aimed to synthesize studies investigating the acute effects of complex whole-body movements with increased parallel information processing on electrical brain activity. A comprehensive search across five scientific databases resulted in thirteen studies meeting the inclusion criteria. The results showed increased theta and alpha activity in frontal, central and parietal areas in most studies during and after movement. However, in other frequency bands the findings were not consistent. Comparisons between complex movements with varying parallel demands revealed a trend towards higher theta for movements with more parallel actions. Based on a consistent EEG methodology, future research should consider movement complexity not only related to the length of sequences but rather in terms of parallel motor activities as a moderator of brain activity to obtain more consistent results in the context of neural effects of movement exercise.

## 1 Introduction

The acute effects of exercise on brain activity have garnered significant attention in recent decades. Conventional physical exercises in the areas of endurance and strength were primarily in the focus of early studies. Meanwhile, there is an increasing interest in physical exercises with a large number of degrees of freedom that is associated with higher coordinative and thus mental demands (Budde et al., 2008; Pesce, 2012). This shift was partly due to the observation of positive effects of sport on executive functions (Benzing et al., 2016). In this context, the electroencephalography (EEG) offered a promising, non-invasive approach with the potential to measure brain activity and distinguish different degrees of cognitive load depending on movement complexity (Sauseng et al., 2007). The EEG signals are typically differentiated by delta (0.5 – 4 Hz), theta (4 – 8 Hz), alpha (8 – 13 Hz), beta (13 – 30 Hz), and gamma frequency bands (>30 Hz; Teplan, 2002). In some studies, a further distinction is made within the alpha and beta bands into alpha-1 (8 – 10.5 Hz), alpha-2 (10.5 – 13 Hz), beta-1 (13 – 15 Hz), and beta-2 (15 – 30 Hz) (Büchel et al., 2021; Hottenrott et al., 2013).

Delta waves are conventionally associated with deep sleep and so far are rarely measured in the context of sport movements (Abhang et al., 2016). However, recent studies also show their involvement in cognitive processing (Harmony, 2013; Malik and Amin, 2017). Theta waves are typically observed during relaxed wakefulness and reduced vigilance (Mari-Acevedo et al., 2019; Zschocke et al., 2012). They were also associated with memory processes (Herweg et al., 2020), the encoding of new information (Klimesch, 1999) and particularly, characterized by frontal midline theta, with attention and executive control processes (Aftanas and Golocheikine, 2001; Cavanagh and Frank, 2014; Cavanagh and Shackman, 2015; Doppelmayr et al., 2008). Sauseng et al. (2007) found a dependence of the anterior midline theta power on the level of mental effort during movements whose complexity was defined by the length of the key press or path sequences. Alpha waves are the dominant waves in the normal waking state. They are also associated with attentional processes and play a role in various memory processes, such as working (Palva and Palva, 2007), semantic (Klimesch, 1997) and long-term memory (Klimesch, 1999). More recent theories have suggested that alpha waves may be related to the active inhibition of task-irrelevant areas (Klimesch et al., 2007). A further differentiation assigns attention processes to the alpha-1 and memory processes primarily to the alpha-2 band (Klimesch, 1999, 1997). Beta waves are associated with anxious thinking, problem solving, and deep concentration (Kirschbaum, 2008; Malik and Amin, 2017). Engel and Fries (2010) attribute beta waves to involvement in attention-related top-down processes and the sensorimotor system. From a more differentiated perspective, the lower beta-1 waves are increasingly evident in focused concentration (Kirschbaum, 2008), while beta-2 waves are characteristic of increased arousal such as stress or excitement (Abhang et al., 2016). Gamma waves play an important role in attention and working and long-term memory (Herrmann and Mecklinger, 2001; Jensen et al., 2007). Additionally, they are participating during the control of connectivity between different brain regions that are crucial for perception, movement, memory, and emotions (Guan et al., 2022).

The interpretation of the frequency bands is based on an assignment of the individual frequency bands to certain activities, usually relying on correlations between activities with simultaneous derivation of brain activity. Since brain activity only ever reflects momentary activities and specific conditions, the interpretations of the frequency bands are always limited to these and cannot be generalized epistemologically. To date, the most common frequency band interpretations have been based almost exclusively on activities recorded in laboratory settings during stationary, seated or lying activities with dominant cognitive tasks.

In recent decades, however, the influence of physical activity on brain oscillations has become increasingly important (Boutcher and Landers, 1988; Kubitz and Landers, 1993). Previous studies have primarily focused on endurance exercises such as running or cycling (for a review see Crabbe and Dishman, 2004; Gramkow et al., 2020; Hosang et al., 2022). Recent advances in data pre-processing and technical developments (such as wireless EEG) enable measurements to be taken during movement (Mehta and Parasuraman, 2013), allowing for more recordings during exercises that involve a larger range of motion. These advances combined with findings from behavioural psychology (Sibley and Etnier, 2003; Tomporowski, 2003) shifted the focus towards a greater emphasis on the mental load caused by a whole-body movement (Baumeister et al., 2008; Budde et al., 2008; Heilmann et al., 2022), which can vary depending on the amount of information that needs to be processed in parallel. The coordinative character of a movement, adaptations to a changing environment, or reactions to opponents and unpredictable situations can increase that kind of information. In the following, exercises that vary in these influencing factors are described as complex exercises. These factors appear to have varying effects on neuronal processes, depending on the complexity of the movement. Even a slight increase in movement complexity seems capable of leading to a neural adaptation. For example, an increased modulation of efficiency in the alpha-network was observed after cross-country skiing compared to conventional running in an laboratory environment. The increased coordinative demand is associated with an increased arm-leg coordination when performing the movement compared to conventional running (Büchel et al., 2023). Previous reviews in the area of movements primarily analyzed endurance exercises with various metabolic loads in combination with EEG, finding the most frequent effects at alpha and beta oscillations, while other frequencies showed inconsistent results (Gramkow et al., 2020; Hosang et al., 2022). However, physical exercises with increased coordinative demand but minimal cardiovascular changes show different results. An increase in theta activity was consistently observed in standing balance exercises (Gebel et al., 2020; Hülsdünker et al., 2015; Wittenberg et al., 2017) or exercises from the field of target shooting training in standing positions (e.g. golf putting, rifle shooting; Doppelmayr et al., 2008; Kao et al., 2013). Due to the growing number of studies investigating complex whole-body movements, a scoping review could give an overview of the effects of complexity of a movement on neural oscillations. Therefore, this review aims to examine studies that analyze cortical activations in healthy individuals triggered by the acute execution of complex movements exceeding the range of motion of pure whole-body balance exercises.

## 2 Method

### 2.1 Study protocol

A scoping review was conducted according to the guidelines of the Preferred Reporting Items for Systematic Reviews and Meta-Analysis (PRISMA) – extension for scoping review (Tricco et al., 2018).

### 2.1 Search strategy, selection process and eligibility criteria

A comprehensive search was conducted on five databases (Pubmed, Web of Science, Scopus, SPORTDiscus and ProQuest) up to 06.06.2024. Appropriate Boolean operators (AND, OR and NOT) were used to join the various keywords. The following term was used for the search: (exercise OR “physical activity”) AND (EEG OR electroencephalography). Duplicated articles were removed using the Endnote software (version 20; The EndNote Team, 2013). The selection process was conducted independently by two authors, and any disagreement between the two authors were solved by consensus. The studies were assessed based on the title and abstract, followed by an analysis of the full text to determine, whether they met the previously defined eligibility criteria. These criteria were: (1) studies had to be peer-reviewed and written in English, (2) the subjects were healthy participants, (3) the intervention had to consist of an acute physical exercise with increased mental demands (complex exercise), (4) studies have made a comparison over a certain measurement time or comparison against another type of sport, (5) studies have measured cortical activity by EEG.

Studies were removed if the exercise consisted of pure endurance (e.g. running, cycling) or strength exercise with a small number of degrees of freedom (e.g. bench press, squats). Pure endurance sports without any mental demand were excluded because of low and monotonous coordinative demand. The aim was to focus mostly on the coordinative aspect. Event-related potential and missing spectral analysis (Büchel et al., 2023; John et al., 2022) led to exclusion as well as studies in which cognitive tests were performed simultaneously with exercise (e.g. dual-task studies; Kahya et al., 2022), as the effects were only of interest triggered by the exercise. Published abstracts or conference papers were also not included in the further analysis. The screening process is shown in Figure 1. Both the search for suitable studies and the quality assessment were carried out independently by the first and second author.

**Figure 1.**
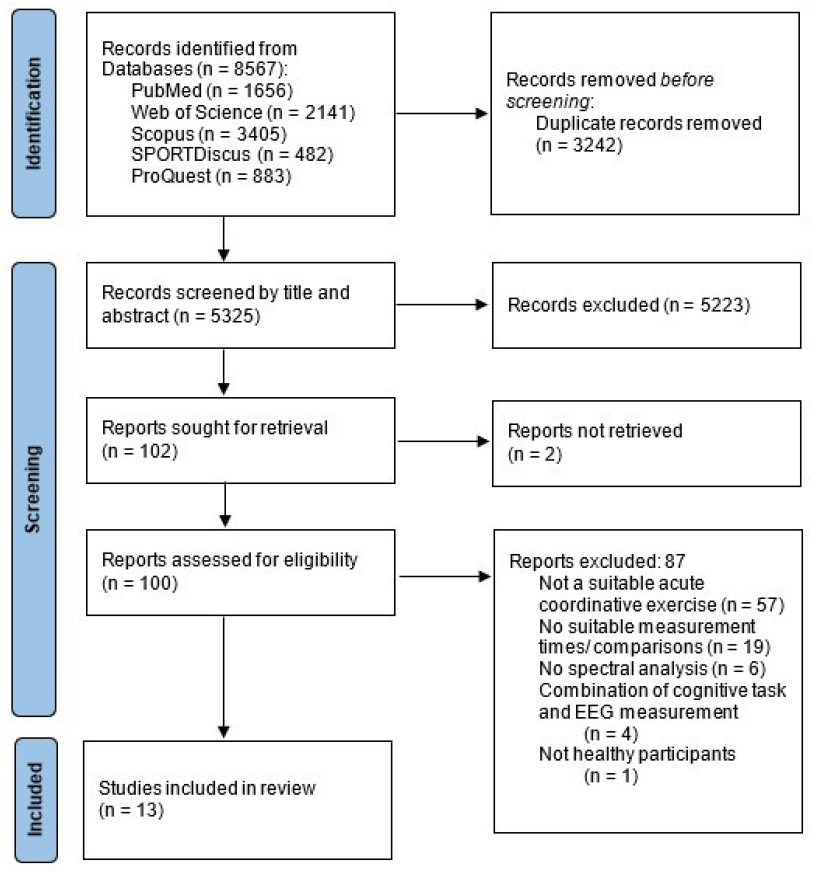
PRISMA flow diagram illustrating the literature search.

**Table 1.**
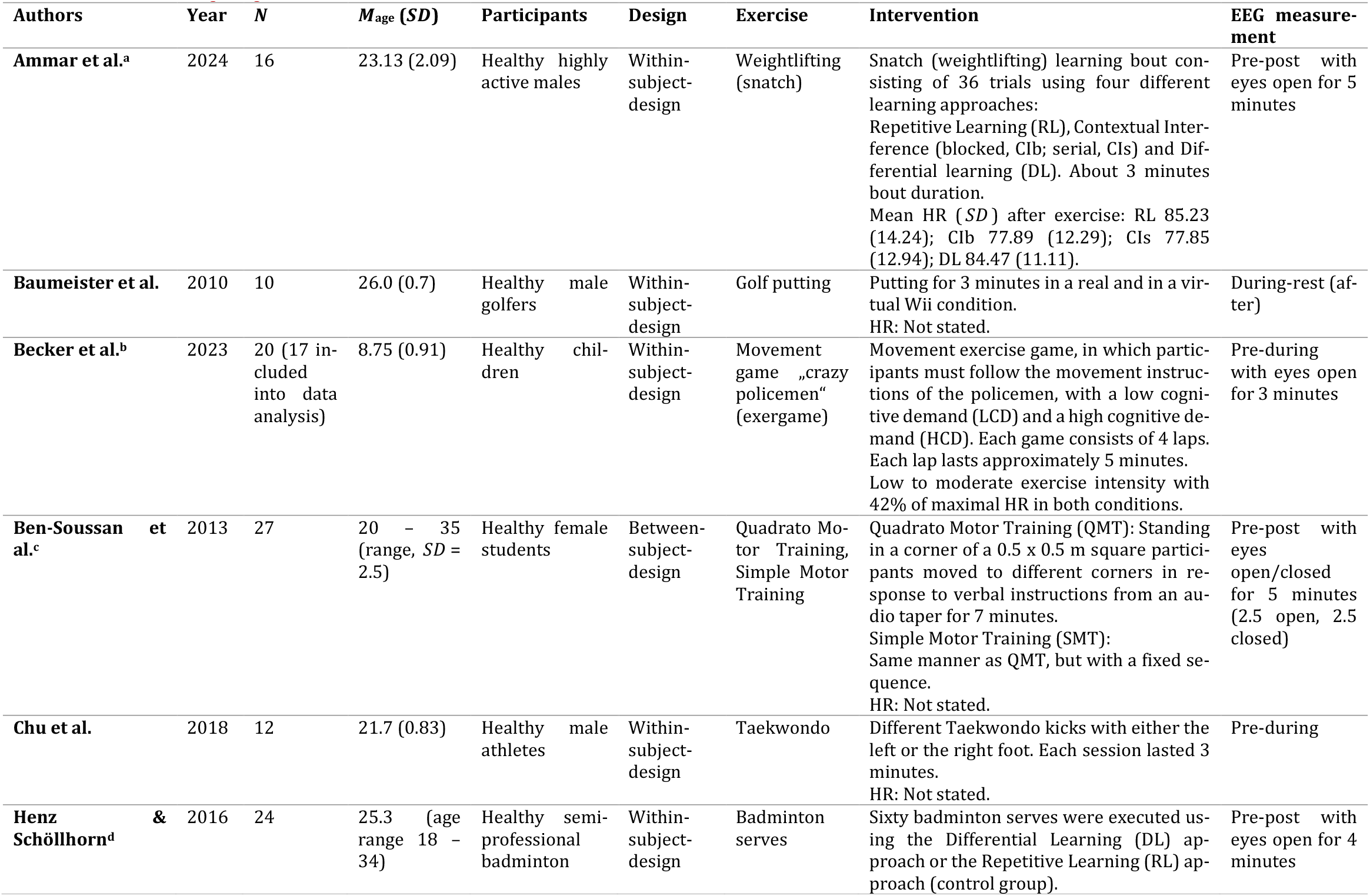
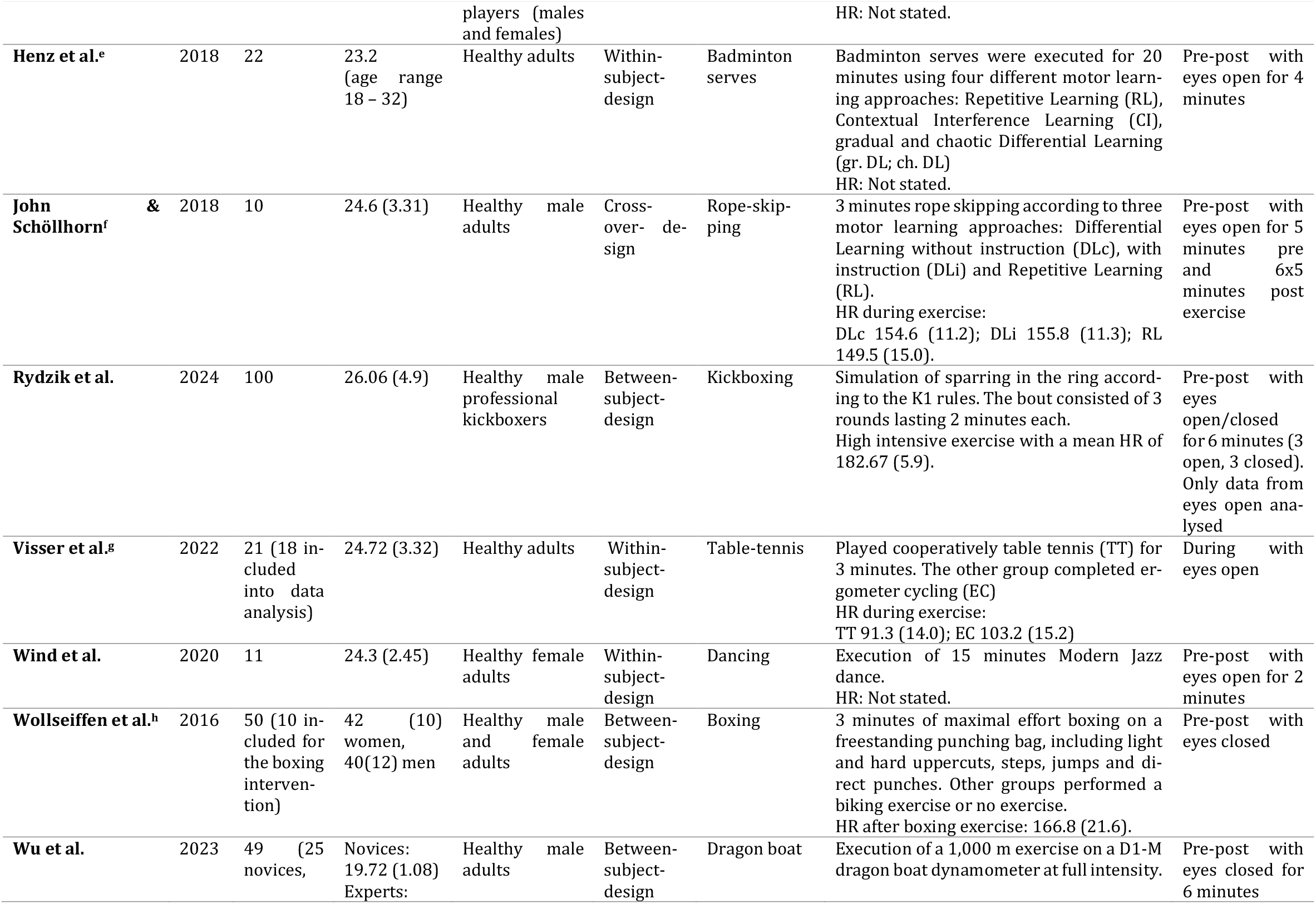
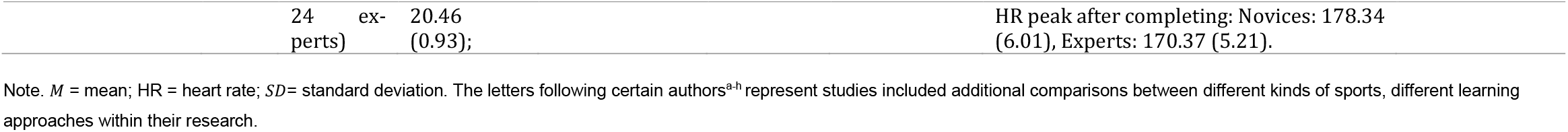
An overview showing design characteristics of the included studies.

**Table 2.**
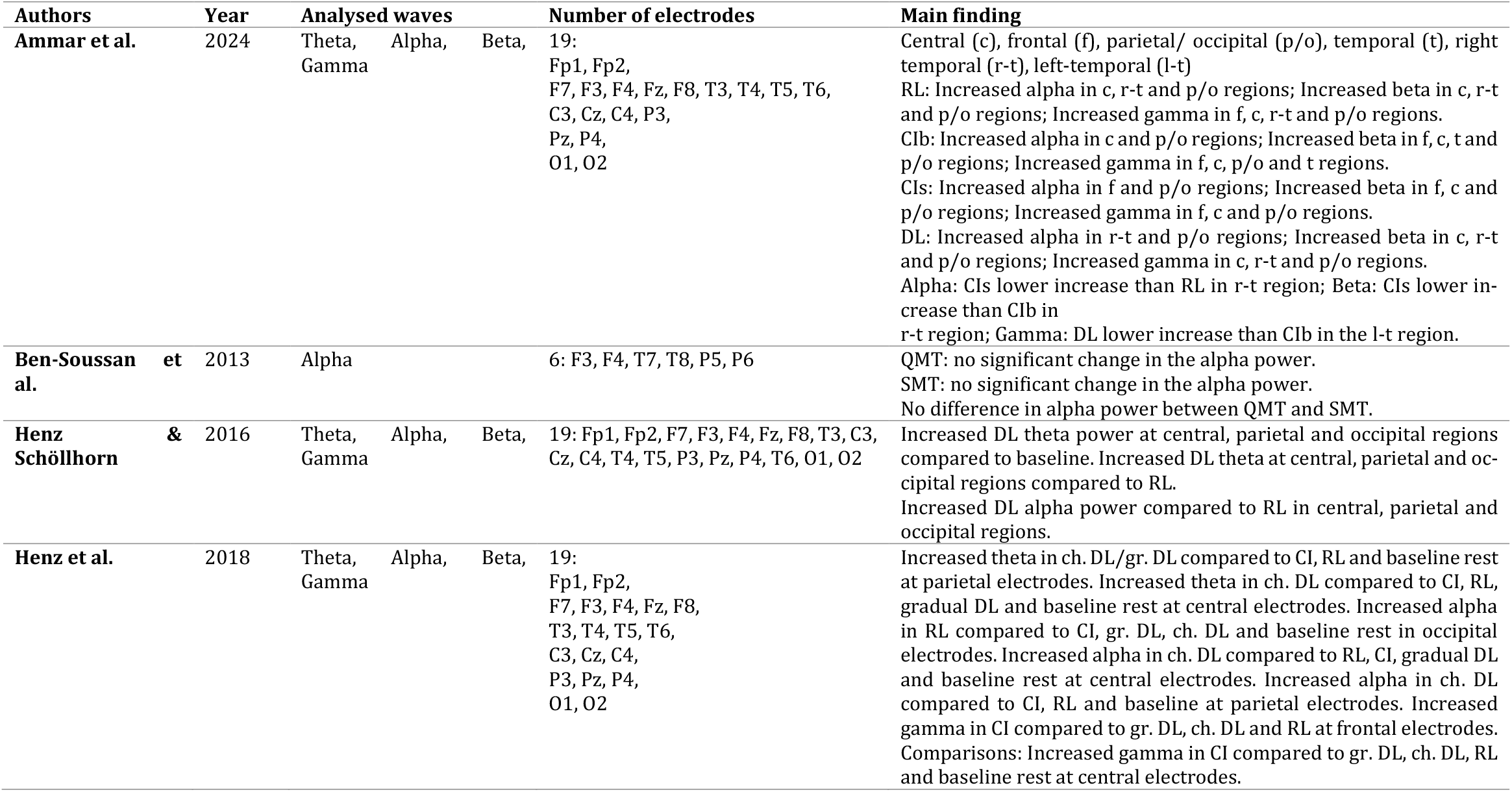
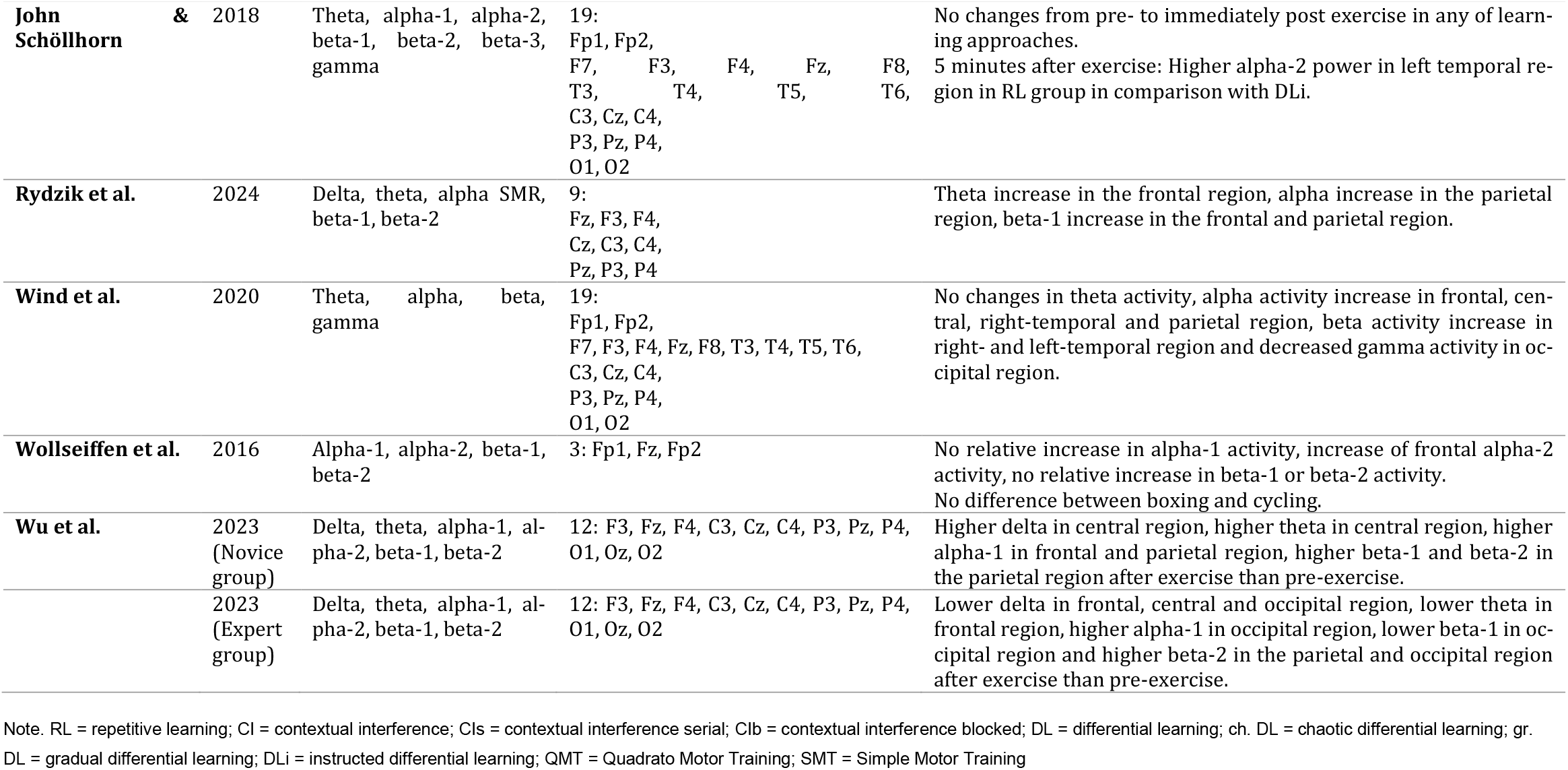
An overview of the EEG-related information included the main findings from pre-post-measurement studies.

**Table 3.** An overview of the EEG-related information included the main findings from during-measurement studies.

| Authors | Year | Analysed waves | Number of electrodes | Main finding |
| --- | --- | --- | --- | --- |
| **Baumeister et al.** | 2010 (Real environment) | Theta, alpha-1, alpha-2 | 6: Fz, F3, F4, Pz, P3, P4 | Significant higher theta in the frontal region, no change in the alpha-1 activity and no change in the alpha-2 activity during the exercise compared to rest. |
|  | 2010 (Virtual environment) | Theta, alpha-1, alpha-2 | 6: Fz, F3, F4, Pz, P3, P4 | Significant higher theta activity in the frontal region, no change in the alpha-1 activity and a significant higher alpha-2 activity in the parietal region during exercise compared to rest. |
| **Becker et al.** | 2023 | Theta, alpha-1, alpha-2 | 64 channels | LCD-condition: Increased theta and alpha-1 activity in the prefrontal region, decreased alpha-1 in left and right motor, left and right parieto-occipital and occipital region, decreased alpha-2 activity in left motor and left parieto-occipital regions during vs. baseline.<br>HCD-condition: Increased theta and alpha-1 in the prefrontal region, decreased alpha-1 in left motor, left and right parieto-occipital and occipital regions, decreased alpha-2 activity in the left motor region<br>LCD vs. HCD: Higher theta in prefrontal region and lower alpha-1 in occipital region in HCD during exercise |
| **Chu et al.** | 2018 | Delta, theta, alpha, beta, gamma | One electrode on the left forehead | Decrease of gamma activity in frontal region with both the left and right foot during exercise compared to before |
| **Visser et al.** | 2022 | Theta | 64 channels | Higher theta power in TT compared to EC in the prefrontal and frontocentral region during the exercise. |

Note. LCD = lower cognitive demand; HCD = higher cognitive demand; TT = table tennis; EC = ergometer cycling.

### 2.3 Data extraction

The author, year, number of participants, average age of participants, participant information, study design, type of exercise, intervention content and intensity of exercise were extracted as important information from each eligible study. To provide an adequate overview of the EEG measurements, information of the examined waves, the length of the measurement, the number of electrodes and the main results were also extracted. This information is presented in Tables 1 – 3.

### 2.4 Quality assessment

The study quality was assessed using a combination of the Quality Assessment Tool for Quantitative Studies (QATQS; “National Collaborating Centre for Methods and Tools,” 2008) and a modified^1^ quality assessment tool to evaluate EEG data acquisition and analysis based on Parr et al. (2021). QATQS includes the six criteria (1) selection bias, (2) study design, (3) confounders, (4) blinding, (5) data collection method and (6) withdrawals and dropouts and is used for the general assessment of study quality. The modified quality assessment tool by Parr et al. (2021) includes the criteria (a) artifact handling, (b) brain wave definition^2^, (c), regional specificity, (d) temporal precision and (e) controls to volume conduction and is used to assess EEG measurements. Each of these 11 criteria was given a quality score of 1 – 3 (1 = strong quality; 2 = moderate quality; 3 = weak quality). As the assessment tool of Parr et al. (2021) is used to assess the EEG measurement, this output is used to assess the (5) data collection method criterion of QATQS. Consequently, for the modified tool of Parr et al. (2021), no weak assessment in criteria a – e leads to a 1 (strong quality) in the criterion data collection method, one weak assessment leads to a 2 (moderate quality) and more than one weak assessment leads to a 3 (weak quality). Overall, a study is assigned the value 1 if it does not show the value weak quality in any of the criteria (1 – 6), a 2 if it shows the value once and a 3 if it shows the value weak quality more than once. A detailed list of the quality assessments is available in Supplementary Table 1.

## 3 Results

The initial search in five databases yielded 8567 records. A total of 3242 articles were removed as duplicates. 5325 articles were screened by title and abstract, of which 5223 studies were excluded by title and abstract. Three studies were not retrievable. One of them was sent to the author upon request. After a careful review of 100 articles, 13 articles were met for the eligibility criteria and included in our scoping review. Further studies were excluded for the following reasons: studies whose exercise was not acute and coordinative according to the above mentioned characteristics (n = 57), no suitable measurement time points or comparisons were made (n = 19), no spectral analysis was performed (n = 6), a cognitive task was solved simultaneously during the EEG measurement (n = 4) or the study did not involve healthy participants (n = 1). This resulted in 13 studies that were included in this review. These can be divided into two groups: Studies that focused on EEG comparison before and after exercise (pre-post, n = 9) and studies that focused on measurement during exercise (during, n = 4).

### 3.1 Complex whole-body exercises

All selected studies (Table 1 – 3) involve an acute complex exercise with healthy participants as intervention. In 12 of the 13 studies, the subjects were adults, while in one study the subjects were children. In terms of content, the sports intervention exercises can be classified as racket sports: badminton (Henz et al., 2018; Henz and Schöllhorn, 2016), golf (Baumeister et al., 2010) and table tennis (Visser et al., 2022); in the area of coordinative demands combined with force-velocity requirements: kickboxing (Rydzik et al., 2024), Taekwondo (Chu et al., 2018), boxing (Wollseiffen et al., 2016) and weightlifting (Ammar et al., 2024); in the area of coordinative demands combined with some endurance requirements: Quadrato Motor Training (QMT, Ben-Soussan et al., 2013), rope skipping (John and Schöllhorn, 2018), dragon boat (Wu et al., 2023); in exergames (“crazy policemen”, with its two variations lower cognitive demand (LCD) and higher cognitive demand (HCD); Becker et al., 2023) and dance (Wind et al., 2020). In four studies, the movements were executed within the framework of various learning models, including repetition learning (RL), contextual interference learning (CI) with its modifications serial (CIs) and blocked (CIb) and differential learning (DL) with its modifications gradual (gr. DL), chaotic (ch. DL), instructed (DLi) and non-instructed (DLc) differential learning. The physical exertion varied between the studies and covered the spectrum from low metabolism related exertion (putting; Baumeister et al., 2010) to maximum intensity of metabolism (dragon boat; Wu et al., 2023). This was measured by the heart rate (n = 7), while in some of the studies no information was provided regarding the metabolism related intensity of the exercise (n = 6).

### 3.2 Studies which compared EEG pre - and post activity

#### 3.2.1. Theta activity

Of the nine studies that measured the effects of exercise on brain activity in a pre-post-test design, seven analyzed the theta band. Four showed statistically significant increases in the frontal region (n = 1; kickboxing, Rydzik et al., 2024), central region (n = 3; e.g. dragon boat (novice), Wu et al., 2023), in the occipital region for badminton serves (n = 1; Henz and Schöllhorn, 2016) and in the parietal region for the same movement (n = 2; ch. DL, gr. DL, Henz et al., 2018; Henz and Schöllhorn, 2016). Three studies found no statistically significant changes (e.g. dance, Wind et al., 2020), while one study reported a significant decrease in theta activity in the frontal region in the context of dragon boat race (expert, Wu et al., 2023).

#### 3.2.2 Alpha activity

All nine studies investigated changes in alpha activity. Six showed a significant increase after exercise compared to before. A further examination of the cortical areas revealed an increase in the frontal (n = 4; e.g. dance, Wind et al., 2020), central (n = 3; e.g. badminton (ch. DL), Henz et al.,

2018), parietal (n = 4; e.g. dragon boat (novice), Wu et al., 2023), occipital (n = 2; RL, Henz et al., 2018; expert, Wu et al., 2023), right temporal (n = 2; RL, DL, Ammar et al., 2024; Wind et al., 2020) and in the parietal/occipital cortex in the context of the snatch (n = 1; Ammar et al., 2024). Wollseiffen et al. (2016) observed an significant increase of alpha activity only in the alpha-2 band after boxing, whereas Wu et al. (2023) reported an enhancement only in the alpha-1 band in both groups of the dragon boat. Three studies (e.g. QMT, Ben-Soussan et al., 2013) reported no changes in the examined cortical areas after the exercise and none of the studies found a decrease in the alpha activity.

#### 3.2.3 Beta activity

Eight of the nine studies analyzed changes in beta activity from before to after activity. Four reported a significant increase in beta activity. These increases were observed in the frontal (n = 2; snatch (CIs, CIb), Ammar et al., 2024; kickboxing, Rydzik et al., 2024), parietal (n = 2; Rydzik et al., 2024; dragon boat, Wu et al., 2023), parietal/occipital (n = 1; Ammar et al., 2024), central (n = 1; Ammar et al., 2024), left temporal (n =2; CIb, weight lifting; Ammar et al., 2024; dance, Wind et al., 2020), right temporal (n = 2; RL, DL, CIb, weight lifting, Ammar et al., 2024; Wind et al., 2020) and occipital region (n = 1; dragon boat, expert, Wu et al., 2023). Rydzik et al. (2024) found an enhancement in the parietal and frontal cortex only in the beta-1 band after kickboxing and Wu et al. (2023) observed the parietal and occipital increase after dragon boat in the expert group only in beta-2 band. Four studies found no changes in beta activity in the examined areas (n = 4; e.g. rope skipping, John and Schöllhorn, 2018). Only one study (Wu et al., 2023), reported a decrease in beta-1 activity in the occipital cortex of the expert group in the context of dragon boat racing.

#### 3.2.4 Gamma activity

Five studies investigated gamma activity. Two reported an increase in gamma activity. This statistically significant increase was observed in the frontal (n = 1; snatch (RL, CIs, CIb), Ammar et al., 2024), central (n = 2; Ammar et al., 2024; badminton (CI), Henz et al., 2018), right temporal (n = 1; RL, DL, CIb, Ammar et al., 2024), left temporal (n =1; CIb, Ammar et al., 2024) and parietal/occipital region (n = 1; Ammar et al., 2024). Henz et al. (2016) as well as John and Schöllhorn (2018) reported no significant change in gamma activity after badminton serves or rope skipping and one study showed a decrease in gamma activity in the occipital region (dance, Wind et al., 2020).

#### 3.2.5 Delta activity

Delta activity was investigated in only two of the nine studies. Wu et al. (2023) observed an increase in delta activity in the central cortex of the novice group in the context of dragon boat racing, but a decrease in the frontal, central, and occipital region of the expert group. However, Rydzik et al. (2024) found no significant changes in delta activity after kickboxing.

The overall quality of the studies that measured before and after the exercise was rated as moderate (M = 2.33).

### 3.3 Studies which measured EEG during the activity

#### 3.3.1 Theta activity

Of the four studies that measured theta activity during movement, all investigated theta activity. Three of the four studies reported an increase during exercise. This increase was observed in the prefrontal cortex during the exergame and table tennis (n = 2; LCD, HCD, Becker et al., 2023; Visser et al., 2022), in the frontal cortex during golf putting compared to after (n = 1; Baumeister et al., 2010) and in the frontocentral region (n = 1; table tennis, Visser et al., 2022). Chu et al. (2018) reported no statistically significant change in theta activity during Taekwondo compared to resting state, and none of the studies showed a decrease.

#### 3.3.2 Alpha activity

Alpha activity was analyzed in three studies. Significant increases were reported in two investigations, namely during golf putting and the exergame (n = 2; Baumeister et al., 2010; Becker et al., 2023). These increases were observed as enhanced alpha-1 power in the prefrontal region (n =1; exergame (LCD, HCD), Becker et al., 2023) and enhanced alpha-2 activity in the parietal region (n =1; virtual golf putting, Baumeister et al., 2010). Chu et al. (2018) reported no changes compared to resting state in alpha activity in Taekwondo movements. One study found a decrease in alpha activity (Becker et al., 2023), specifically a decrease in alpha-1 power in the left and right parieto-occipital region (LCD, HCD), occipital region (LCD, HCD), left motor cortex (LCD, HCD), and right motor cortex (LCD) during the exergame compared to before. Additionally, they showed a reduction in alpha-2 power in the left parieto-occipital region (LCD) and left motor cortex (LCD, HCD).

#### 3.3.3 Beta activity

One study investigated the beta activity during exercise (Chu et al., 2018). The authors reported no significant change of beta activity compared to before.

#### 3.3.4 Gamma activity

Gamma activity was also investigated in only one study (Taekwondo, Chu et al., 2018). This study showed a significant decrease in gamma activity in the frontal region.

#### 3.3.5 Delta activity

Delta activity was also investigated in only one study (Chu et al., 2018). This study showed no significant change in activity during the exercise compared to before.

The overall study quality in this area was rated as moderate (M = 2.5).

### 3.4 Studies comparing exercises

In eight^a-h^ of the 13 studies, a further comparison was made between exercises that differed in their coordinative demands from the intervention exercise (see letters in Table 1). This comparison was made in two studies on cycling (table tennis, Visser et al., 2022; boxing, Wollseiffen et al., 2016), in four studies the exercise was performed using different learning approaches with different variabilities (snatch, Ammar et al., 2024; badminton serve, Henz et al., 2018; badminton serve, Henz and Schöllhorn, 2016; rope skipping, John and Schöllhorn, 2018) and in two studies the movement was varied by externally specifying the direction of movement and thus requiring a situational response with increased spatial orientation (exergame, Becker et al., 2023; QMT, Ben-Soussan et al., 2013). Wollseiffen et al. (2016) and Ammar et al. (2024) compared the increases from pre to post, while the others focused on the differences in the absolute EEG power between the movements at the respective measurement time.

#### 3.4.1 Theta activity

Of the eight studies that investigated differences in frequencies between exercises, six^abdefg^ studies investigated theta activity. Four reported differences in theta activity between exercises. This was reflected in an increase in the prefrontal (n = 2; HCD^3^, Becker et al., 2023; TT, Visser et al., 2022), frontocentral (n = 1; TT, Visser et al., 2022), central (n =2; chaotic DL, Henz et al., 2018; DL, Henz and Schöllhorn, 2016), parietal (n = 2; chaotic DL, Henz et al., 2018; DL, Henz and Schöllhorn, 2016) and occipital cortex (n = 1; DL, Henz and Schöllhorn, 2016). Two studies that executed the movements of snatching and rope skipping using different movement learning models found no significant differences in theta activity between the movement sequences (Ammar et al., 2024; John and Schöllhorn, 2018).

#### 3.4.2 Alpha activity

Alpha activity was investigated in seven^abcdefh^ of the eight studies. A difference in alpha activity was reported in five studies. This was found in the right temporal (n = 1; snatch (RL), Ammar et al., 2024), left temporal (n = 1; rope skipping (RL), John and Schöllhorn, 2018), occipital (n = 3; e.g. LCD, Becker et al., 2023), parietal (n = 2; e.g. chaotic DL, Henz et al., 2018) and central region (n = 2; chaotic DL, Henz et al., 2018; DL, Henz and Schöllhorn, 2016). The difference by Becker et al. (2023) in the occipital region was only shown in alpha-1 activity and the difference in the left temporal region by John and Schöllhorn (2018) was only shown in alpha-2 activity. Two studies reported no difference in alpha activity between exercises (Ben-Soussan et al., 2013; Wollseiffen et al., 2016).

#### 3.4.3 Beta activity

Beta activity was examined in only five^adefh^ of the eight studies. Only Ammar et al. (2024) showed a greater increase in the group with a blocked sequence of exercise (Contextual interference blocked) compared to the Contextual interference serial group in the right temporal cortex. Four of the remaining studies showed no significant difference between the groups (n = 4; e.g. Wollseiffen et al., 2016) during or after intervention.

#### 3.4.4 Gamma activity

Four^adef^ of the eight studies investigated gamma activity. Two studies showed differences between the exercises. These were found in left temporal (n = 1; CIb, Ammar et al., 2024), frontal (n = 1; CI, Henz et al., 2018) and central cortex (n = 1; CI, Henz et al., 2018). Two studies reported no differences in gamma activity (Henz and Schöllhorn, 2016; John and Schöllhorn, 2018).

#### 3.4.5 Delta activity

None of the studies listed here examined delta activity.

The overall quality of the studies, which included several exercises with different coordinative requirements, was rated as moderate (M = 2.25).

## 4 Discussion

The aim of this review was to investigate the neural effects of acute mental demanding physical exercises. This demand is typically found in complex movements involving many degrees of freedom in parallel. The neural effects were investigated by spectral EEG activity either in a pre-post-test design or in a during design. In some of the studies, a further comparison was made between exercises of different coordinative demands. After the selection process, there were 13 studies left (pre-post = 9; during = 4). In 12 of the 13 studies the participants were healthy adults, in one study healthy children. The duration of the physical activities ranged from 3 to 20 minutes. Both cyclical (e.g. Wu et al., 2023) and noncyclical (e.g. Henz and Schöllhorn, 2016) movements were analyzed.

### 4.1 Effects on theta frequency

Seven of the eleven (64%, Figure 2) studies that examined theta frequency reported significant increases (pre-post-design = 4; during-design = 3). Three studies showed no change, and one reported a decrease. Increases were found in the central (n = 3; e.g. Henz and Schöllhorn, 2016), frontal (n = 2; Baumeister et al., 2010; Rydzik et al., 2024), prefrontal (n = 2; Becker et al., 2023; Visser et al., 2022) and frontocentral regions (n = 1; Visser et al., 2022). Theta activity in primary frontal areas is linked to cognitive tasks requiring increased executive control (Cavanagh and Frank, 2014; Cavanagh and Shackman, 2015; Jensen and Tesche, 2002) and attention (Aftanas and Golocheikine, 2001; Gevins, 1997). Similar findings have been observed in exercises in the context of sports. For example, increased frontal midline theta values were found in targeting exercises in the pre shooting phase or balance exercises that required an increased level of executive control (Doppelmayr et al., 2008; Hülsdünker et al., 2016; Sipp et al., 2013). However, these increases could not be shown for cyclical, mostly automated movements with less coordinative requirements (Gramkow et al., 2020; Hosang et al., 2022). A somewhat more specific view shows that all studies that examined movements from the area of racket sports (n = 4; e.g. Baumeister et al., 2010) showed an increase in theta frequency. This may be attributed to the specific requirements of the movement, which are not only characterized by higher demands on coordination and attention processes, but e.g. in the case of table tennis also by the additional situational reactions to the opponent, which increase the information to be processed by the visual information input and to which the corresponding reactions must be adapted (Visser et al., 2022). This additional visual information means that information from the visual cortex must be integrated into the whole-body sensory and motoric activities. The synchronization of areas that are more distant tends to be attributed to lower frequencies such as theta frequencies (Varela 2001; Fries 2005). Since similar effects were found with balance exercises (Gebel et al., 2020; Hülsdünker et al., 2015) for example, where multiple joints, visual and vestibular information have to be integrated in parallel to keep the centre of gravity within the support area, common movement requirements as increased cognitive control especially in frontal areas (Cavanagh and Frank, 2014) could be responsible. When considering single frequency bands and only the type of sports in general, the remaining categories of the examined complex exercises show inconsistent results and give reason to further investigate the movement-specific demands. One reason, that not all studies showed an increase could lie in the factors influencing EEG activity, as reported in the study by Wu et al. (2023), which identified higher theta activity in novices than in experts in the context of dragon boat race, indicating a shift of focus with increasing expertise.

**Figure 2.**
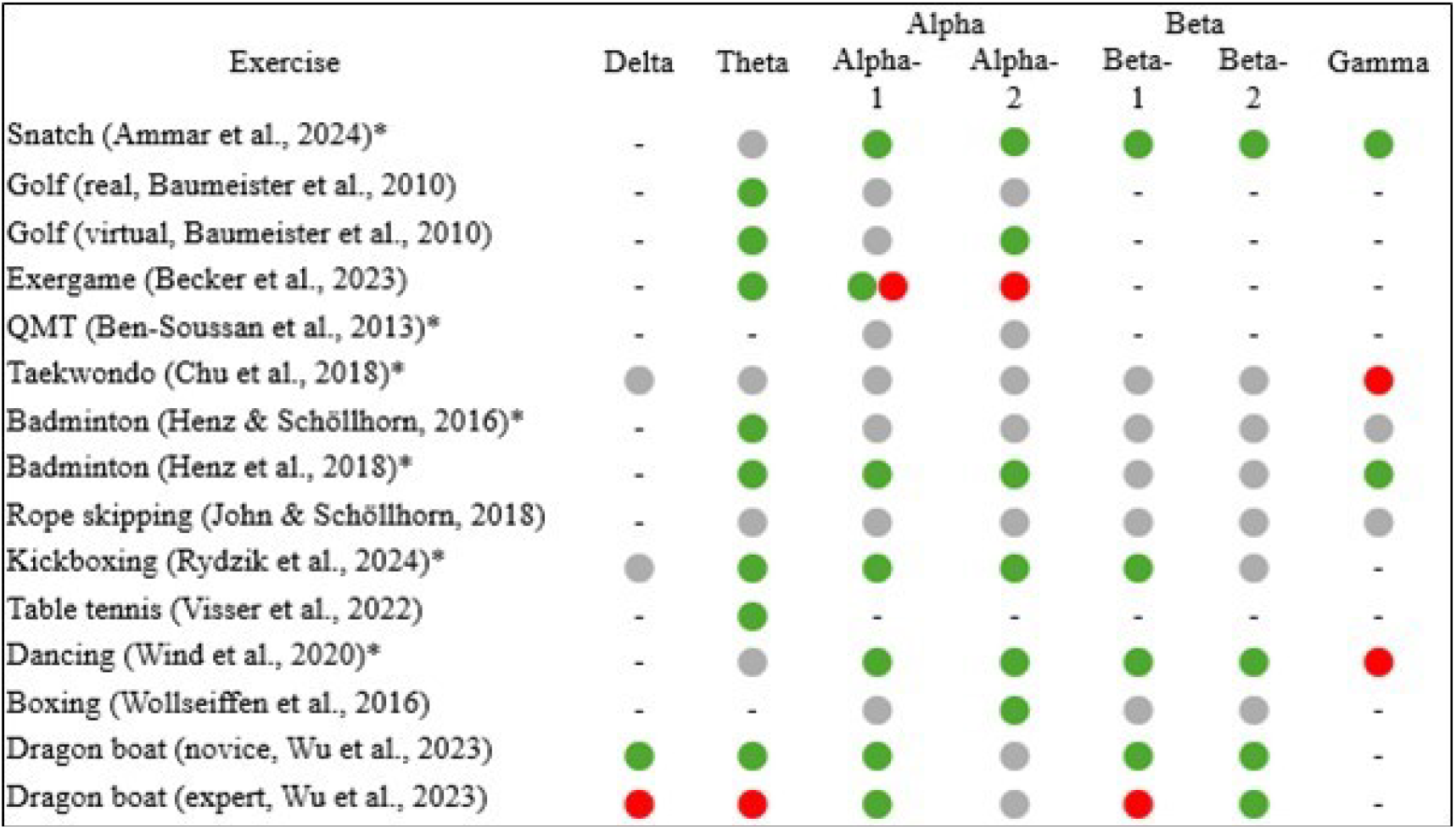
Overview of frequency-band specific EEG results from pre-post measurements and pre/after-during measurements. Green indicates an increase, red a decrease, a combination of both an increase and a decrease, grey indicates no change. A bar (-) indicates that the frequency was not analyzed. An asterisk after the author indicates that no distinction was made between alpha-1 und alpha-2 or beta-1 and beta-2 (Rydzik et al. (2024) only distinguished between beta-1 und beta-2), QMT = Quadrato Motor Training.

Some of the studies carried out a further analysis by comparing movements of different mental demands. This examined whether the neural activities differed between the movements. Four^bdeg^ of the six studies that examined theta reported higher values in favour of the movement that had, in comparison, a higher complexity due to a higher coordinative demand. Visser et al. (2022) showed this by comparing a table tennis exercise with an exercise on a bicycle ergometer, and Henz and Schöllhorn (2016) performed badminton serves using different learning approaches with increasing variations within the movement. These comparisons show, on the one hand, that different movements can have varying effects on frontal theta, and on the other hand, that a variation within the same movement can influence this as well and thus confirms the suggestion of Sauseng et al. (2007), that the frontal theta is related to the amount of mental effort. However, these results were not observed in all comparisons.

Four studies found no significant theta changes triggered by exercise (e.g. rope skipping, John and Schöllhorn, 2018). One of the reasons for the differences in the results could be the different demands of the movement on cognition or somatosensory execution. Ermutlu et al. (2015) found differences in the rest theta activity of fast ball sports athletes and dancers, demonstrating the divergent effect of sport on neural activity. Additionally, an interaction effect between metabolic effects and mental demand on theta activity cannot be excluded at this point either and requires further investigation.

Overall, the majority of studies showed theta increases in frontal and central areas. This differs from previous reviews (Crabbe and Dishman, 2004; Gramkow et al., 2020; Hosang et al., 2022), which reported inconsistent results with regard to theta activity in endurance dominated exercises. The results presented here are consistent with findings from the field of shooting (Doppelmayr et al., 2008) or balancing (Hülsdünker et al., 2016) and extend the findings that similar effects on theta activity can also be found in exercises with an increased duration or intensity where the influence of various fatigue mechanisms come in additionally. Here rather more differentiation is recommended instead of hasty generalizations. In addition, the further comparisons between the exercises provide indications of a correlation between the complexity of the movement and theta activity, which should be pursued further.

### 4.2 Effects on alpha frequency

Alpha activity was examined in 12 of 13 studies. Eight (67%) of these reported significant increases (pre-post-design = 6; during-design = 2), four studies showed no significant changes, and one reported a decrease. The increases were primarily observed in frontal (n = 5; e.g. Wollseiffen et al., 2016), central (n = 3; e.g. Wind et al., 2020) and parietal regions (n = 6; e.g. Rydzik et al., 2024). Our results are partially consistent with previous research, which also found significant alpha increases during and after endurance dominated exercises in frontal and central areas of the brain (Hosang et al., 2022). Hosang et al. (2022) argued that this was due to the inclusion of cognitive functions during and after exercise. Frontal alpha synchrony is associated with top-down control and inhibition processes when measured during activities (Klimesch et al., 2007; Misselhorn et al., 2019) and thus offers an explanation for the increase in these areas. However, the approach hardly explains the increase in alpha power when eyes are closed and why children under five years of age have this as highest frequency. The increased alpha activity in the parietal cortex differs from previous research findings. Parietal alpha activity is typically associated with spatial orientation (van Schouwenburg et al., 2017), internal attention (Benedek et al., 2014) and intersensory reorientation (Misselhorn et al., 2019). All three associations are typically considered as a necessity for whole-body athletic movements.

The simultaneous involvement of frontal and parietal areas is consistent with previous hypothesis that the frontoparietal network is involved in the top-down control of attention (Corbetta and Shulman, 2002; Noudoost et al., 2010). While frontal lobe activity is more associated with top-down control, parietal activity is rather assigned to bottom-up control (Buschman and Miller, 2007). Regarding the increased demands on various attentional processes to successfully execute the exercises investigated here, this can be used as a possible explanation for the increased parietal alpha activity. Of the reported increases in alpha activity, two studies (Becker et al., 2023; Wu et al., 2023) only reported changes in alpha-1 frequency and two (Baumeister et al., 2010; Wollseiffen et al., 2016) only showed these in alpha-2 activity.

Seven studies further compared different movements, of which five^abdef^ studies reported differences in alpha activity. However, this extended to different parts of the cortex and thus does not show uniform activation in a specific area. Only three studies reported significant differences in the occipital cortex (e.g. Becker et al., 2023), which is associated with both visual processes (Zschocke et al., 2012) and working memory (Tuladhar et al., 2007). These differences may indicate different demands of movement on visual or working memory requirements.

Four studies reported no differences triggered by the exercise (e.g. Chu et al., 2018), whereas Becker et al. (2023) reported a significant decrease during a movement exergame in alpha activity. This exercise was performed, analogous to the virtual condition of Baumeister et al. (2010), in the context of an exergame and effects due to the digital presentation should not be neglected (Anders et al., 2018; Müller et al., 2023).

In summary, the majority of studies reported alpha increases. These were primarily identified in frontal, central, and parietal areas. The increases in the frontal and central areas are consistent with the findings of previous research on EEG studies with a dominant endurance component. The parietal increase could also be a response to coordinative demands in connection with spatial orientation, as these were also found in cognitive tasks that specifically required spatial attention, internal attention and intersensory reorientation (Benedek et al., 2014; Misselhorn et al., 2019; van Schouwenburg et al., 2017). Even though the alpha activity is increasingly shown in the complex movements investigated here, the coordinative demand cannot be considered solely responsible for the increase. However, there are tendencies that the coordinative aspect has a positive effect, which is shown in an additional parietal increase. These activities occur across different intensities and exercise durations.

### 4.3 Effects on beta frequency

Beta activity was examined in nine of the 13 studies. Four studies (44%) reported significant increases in activity triggered by sport (pre-post-design = 4), five found no changes, and one study showed a decrease in beta activity (beta-1 in expert, Wu et al., 2023). The increases were shown only sporadically in the temporal, central, and frontal regions, but increasingly in the parietal and occipital cortex (n = 3; e.g. Rydzik et al., 2024). Among the studies that reported significant increases, exercises with larger strength or speed participation as well as increased endurance requirements were represented.

Beta activity is typically associated with visual perception (Piantoni et al., 2010) or working memory (Deiber et al., 2007). Hereby it is important to keep in mind that this assignment is based on the working memory related to Baddeley and Hitch’s (1974) introduction that is related to sequential, visual-spatial tasks and should not be generalized to proprioceptive, kinesthetic and tactile tasks (Baddeley, 2018). Beta oscillations are often observed in various cortical areas, including the occipital (visual attention; Gola et al., 2013) or parietal area (visual-spatial-working memory; Deiber et al., 2007). In the context of movement execution, beta is increasingly reported in sensorimotor processes (Engel and Fries, 2010; Lalo et al., 2007). However, it is difficult to separate sensorimotor control and attention especially in sports, as sensorimotor behaviour is often accompanied by attentional processes (Engel and Fries, 2010). Therefore, the increases found in four studies may be the result of sensorimotor processes, increased attentional demands or working memory processes, which have been shown primarily in the occipital and parietal areas due to the specific cognitive demands, but less pronounced in the central areas. The assumption that the activation of beta oscillations is not solely the result of sensorimotor movement is strengthened by the fact that beta increases were shown in areas that are not only associated with sensorimotor processes (e.g. Rydzik et al., 2024).

Five^adefh^ of the studies analyzed beta activity between different movements. One study on weightlifting acquisition showed a different activation in the right temporal area when comparing a movement according to different learning approaches (Ammar et al., 2024). The four remaining studies showed no difference and thus no consistency of the results at this frequency.

Five studies reported no significant changes after or during exercise and one indicated a decrease in beta-1 activity in the occipital region (expert, Wu et al., 2023). It should be noted that five of the nine studies did not differentiate beta activity. Given the large range of 13 – 30 Hz, this may have led to missing differences.

In sum, the results for beta activity are not consistent: Four studies found significant increases, five showed no changes and one reported a decrease. Increases were primarily observed in the occipital and parietal cortex and could be attributed to increased demands on somatosensory processes, working memory or visual processes. (Deiber et al., 2007; Engel and Fries, 2010; Gola et al., 2013; Lalo et al., 2007). Our results thus differ regarding beta activity of previous reviews with healthy subjects who have reported frequent increases (Hosang et al., 2022). Whether the differences are caused by different levels of increased metabolism going along with endurance tasks needs further research. However, this difference also could be due to the predominantly shorter exercise duration, the significantly lower average exercise intensity or the increased coordinative demands within the exercises.

### 4.4 Effects on gamma and delta frequency

Gamma oscillations were examined in only six of the 13 studies. Two (33%) reported significant increases (prepost-design = 2), two found no changes, and two showed a significant decrease. While significant increases were shown in the central area (Ammar et al., 2024; Henz et al., 2018), decreases were reported in the occipital and frontal cortex (Chu et al., 2018; Wind et al., 2020). Gamma activity is associated with movement control (Ball et al., 2008; Ulloa, 2022), attention (Jensen et al., 2007) or working memory processes (Thompson et al., 2021). Nevertheless, it is unclear whether gamma oscillations implement causal mechanisms of specific brain functions or represent a dynamic mode of neural circuit function. Therefore, Fernandez-Ruiz et al (2017), speculate that gamma does not represent cognitive activity, but rather an activity motif that describes processes underlying information processing in its brain circuitry. The results presented here are in line with previous findings (Gramkow et al., 2020; Hosang et al., 2022), which also found inconsistent results with regard to gamma activity. Both, the increases and decreases, could be the result of symbiotic effects of cognitive and somatosensory processes, as gamma effects have been shown in both contexts (Ball et al., 2008; Thompson et al., 2021; Ulloa, 2022). Four^adef^ of the studies also compared gamma activation between different exercises or after a variable execution of the movement. Only two studies (Ammar et al., 2024; Henz et al., 2018) reported significant different activations in the context of movement learning models, which, however, did not consistently emerge in a specific region.

Delta oscillations are the least studied frequencies and were only analyzed in three studies (pre-post-design = 2; during-design = 1). Wu et al. (2023) found a significant increase in delta activity in the beginner group and a decrease in the experienced group. No changes were found in two studies (Chu et al., 2018; Rydzik et al., 2024). These differences may be the result of the different performance levels of the respective groups. Lardon and Polich (1996) demonstrated differences in resting EEG delta activity between individuals who engage in regular physical activity and those who are less active. This, in combination with the fact that delta activity is also associated with cognitive processing (Harmony, 2013), could explain the differences in this frequency band, as postural control in dragon boat is thought to require greater cognitive engagement in beginners. However, because of the small number of studies that included delta wave analysis it seems very speculative to assign any of these frequencies to sports movements.

In summary, no consistent conclusion can be drawn for either gamma or delta activity. Both have been investigated in only a few studies and show inconsistent results. Nevertheless, analyzing these oscillations in future investigations would be beneficial, since on the one hand, contrary to previous assumptions, the delta frequency in particular is increasingly associated with cognitive processes (Harmony, 2013) and on the other hand, a basis should be created to be able to analyze potential interaction effects between the frequency bands (Varela et al., 2001).

### 4.5 EEG and its caveats

The studies investigated here show differences in terms of EEG measurement and data processing. The modified criteria by Parr et al. (2021) for the assessment of EEG analyses and the associated effects on study quality (overall moderate study quality; M = 2.38) reveal the importance of the measurement and especially of the correct pre-processing steps. The EEG is an excellent device for recording brain activity, which is, however, contaminated by various external sources. Advanced denoising techniques, such as independent component analysis (ICA), play an important role in this process. Their primary aim is to distinguish brain activity from external recorded activity, such as muscle artifacts (Albera et al., 2012). Among the 13 studies, only seven studies (e.g. Visser et al., 2022) reported using advanced denoising techniques. This is problematic because, in absence of the investigator’s expertise, the frequently reported frontal activity could be contaminated by eye movements or, due to continuous data collection, muscle activity might not be adequately detected and removed manually. Only two of the four studies (Becker et al., 2023; Visser et al., 2022) that measured during movement used techniques such as ICA, which further complicates the interpretation of the data, as movements usually cause increased artifacts. In the study by Chu et al (2018) the pre-processing was carried out automatically by a ThinkGear™ chip, making it impossible to trace the data.

Only five studies (see Figure 2) differentiated alpha activity into alpha-1 and alpha-2 and four differentiated beta activity into beta-1 and beta-2. Increased differentiation would have been beneficial for two reasons. Firstly, a differentiated analysis would have identified potential differences in the subbands and secondly, the relationship between cause and effect could have been demonstrated more effectively, as the subbands are typically associated with different functions. Moreover, the number of electrodes significantly impacts the quality of the data, so that the quality generally increases with the number of electrodes (Lau et al., 2012). The average number of electrodes within the studies was 20, which is sufficient to measure an EEG with good quality (Miraglia et al., 2021). However, two studies utilized a limited number of electrodes. Chu et al. (2018) used only one and Wollseiffen et al. (2016) used three electrodes for their measurements. The fact that none of all studies controlled volume conduction further complicates the spatial interpretation of the locally measured activity (Parr et al., 2021; Rutkove, 2007) This is one of the main reasons for the overall moderate study quality and might be responsible for the inconsistent results across some frequencies. In addition, study designs with mixed genders as well as with athletes and non-athletes were included in this review, which may have further contributed the differing results (Corsi-Cabrera et al., 1993; Fang et al., 2022) The different methods of EEG measurement and pre-processing used in the respective studies make the comparison more difficult and therefore it is pleaded at this point to reach a common consensus regarding the EEG measurements and especially the pre-processing of the data, which is generally recognized. The fact that both the posture of the body (Chang et al., 2011; Jung et al., 2020), and the location during the measurement (indoor/outdoor; Boere et al., 2023) have an influence on the measurement should also be taken into account. The heterogeneity resulting from the different methods is one of the reasons why no quantitative synthesis was carried out in this study.

## 5 Limitations and future research

In general, the limitations of the study are given by the boundary conditions of the study design, and therefore do not allow for generalization nor claim to be comprehensive. Although this review indicates that neural effects may be attributable to coordinative demands, this cannot be definitively concluded, as they may also have increased due to metabolic activity (John et al., 2020). The absence of heart rate measurements in some studies further complicates the assessment of exercise intensity and, consequently, the evaluation of metabolic processes. Furthermore, the analyzed movements are very heterogenous in terms of the complexity of the movement. In particular, the acyclic movements were characterized by pauses between the movements, for which details were sometimes not provided. In this review, brain activity was measured both during and after the movement and mostly compared with activity before the execution. Even if measurements were taken immediately after the movement, the effect may have been reduced in the meantime. In addition, Baumeister et al. (2010) only showed a comparison with activity during and after the movement, so that the activity cannot be clearly compared to activity before the movement.

The years of publication reveal the growing relevance of the topic. In the context of movement related EEG studies it would be helpful for systematic findings to differentiate between changes of brain activation in the context of cognitive demands that are related e.g. to mathematical or language tasks as they are very often used in dual task paradigms (e.g. counting while walking), brain activation caused by increased metabolism (e.g. graded exercise test on cycle ergometer, Gutmann et al., 2018), most often associated with an increase in heart rate or changes of brain activation caused by an increase in complexity resulting in increased information that has to be processed in parallel. New measurement devices allow the application of EEG in another context as well. Future research could investigate other movements or games with different aims. The study by Visser et al. (2022) has shown that measurements taken during an open skill exercise also provide useable information. In this context, it would be interesting to see the neural effects of an increase of the difficulty within the open skill exercise, for example by increasing the pace of play. The influences of both increased intensity and increased coordinative demand would be of interest here. At the same time, the influence of psychological parameters in parallel to complex movements is necessary to prove their generalization from laboratory situations. Measurements immediately after the exercise are also useful in the field of cognitive learning research (Sibley and Etnier, 2003; Tomporowski, 2003) and enable the study of exercises, where measurements would not have been possible during the movement due to the susceptibility of the EEG. As this review only examined exercises lasting a maximum of 20 minutes, it raises the question of how the effects might differ with exercises lasting an hour or more. How do the frequencies, with a particular focus on the frontal and central theta frequency, behave over the time span of an exercise or within the exercise with constantly increasing mental demands? How do pauses between movements influence neural activity. In this research context, potential moderator variables should be considered.

## 6 Conclusion

Technical innovations combined with a shift in the focus of movements have led to an increased investigation of complex whole-body movements with many parallel actions. Based on the main findings of this scoping review, namely a frequently reported increase of theta and alpha activity particularly in frontal, central and parietal areas, there is a need to further research complex movements. Since the number of studies is still small and the heterogeneity of the complex movements examined here is large, more investigations are required to explore the influence of complexity in more detail. Based on a consistent methodological EEG approach, future studies should take the complexity of a movement and the resulting coordinative demands as a potential moderator into account. In view of the beneficial role of the lower frequencies in general learning and therapeutic processes stimulated by the complex movement, the extent of the consequences is only alluded to here, but seems sufficient to intensify future research in this direction.

## 9 Author Contributions

GM and AS conducted the literature search, data collection, and quality assessment. GM wrote the first draft of the manuscript under the supervision of WS. GM and WS have contributed equally to the study design. All authors reviewed and approved the submitted version.

---

**Supplementary Table 1.** Assessment of study quality based on QATQS (National Collaborating Centre for Methods and Tools, 2008) and Parr et al. (2021).

| Study/ author | Se-lection bias | Study De-sign | Con-found-ers | Blind-ing | Data collec-tion method | With-draw-als and drop-outs | Arti-fact han-dling | Defi-nition of brain wave | Re-gional speci-ficity | Tem-poral preci-sion | Con-trols to vol-ume con-duc-tion | To-tal |
| --- | --- | --- | --- | --- | --- | --- | --- | --- | --- | --- | --- | --- |
| Ammar et al. 2024 | 2 | 1 | 2 | 2 | 3 | 1 | 1 | 3 | 1 | 2 | 3 | 2 |
| Baumeister et al. 2010 | 2 | 1 | 2 | 3 | 3 | 1 | 2 | 2 | 3 | 1 | 3 | 3 |
| Becker et al. 2023 | 2 | 1 | 1 | 3 | 2 | 1 | 1 | 2 | 1 | 1 | 3 | 2 |
| Ben-Sous-san et al. 2013 | 2 | 1 | 3 | 3 | 3 | 1 | 2 | 3 | 2 | 2 | 3 | 3 |
| Chu et al. 2018 | 2 | 2 | 3 | 3 | 3 | 1 | 3 | 3 | 3 | 1 | 3 | 3 |
| Henz & Schöllhorn 2016 | 2 | 1 | 2 | 2 | 3 | 1 | 2 | 3 | 1 | 2 | 3 | 2 |
| Henz et al. 2018 | 2 | 1 | 2 | 2 | 3 | 1 | 1 | 3 | 1 | 2 | 3 | 2 |
| John & Schöllhorn 2018 | 2 | 1 | 1 | 3 | 2 | 1 | 1 | 2 | 1 | 2 | 3 | 2 |
| Rydzik et al. 2024 | 2 | 1 | 2 | 3 | 3 | 1 | 3 | 2 | 2 | 2 | 3 | 3 |
| Visser et al. 2022 | 2 | 1 | 1 | 3 | 2 | 1 | 1 | 2 | 1 | 1 | 3 | 2 |
| Wind et al. 2020 | 2 | 1 | 2 | 3 | 2 | 1 | 2 | 3 | 1 | 2 | 2 | 2 |
| Wollseiffen et al. 2016 | 2 | 1 | 2 | 3 | 3 | 1 | 2 | 2 | 3 | 2 | 3 | 3 |
| Wu et al. 2023 | 2 | 2 | 2 | 3 | 2 | 1 | 1 | 2 | 2 | 2 | 3 | 2 |

## Footnotes

1. As this review addressed a different question, one component of the assessment tool of Parr et al. (2021), namely ‘secondary measures of verbal and conscious processing’ was removed as it was not relevant for our purposes.

2. The original component ‘alpha definition’ of Parr et al. (2021) was modified to ‘brain wave definition’ similar to Hosang et al. (2022) because all EEG frequencies were of interest

3. In this section, the abbreviation in parentheses indicates the exercise that exhibits higher EEG value in comparison.

## References

Abhang, P.A., Gawali, B.W., Mehrotra, S.C., 2016. Technical Aspects of Brain Rhythms and Speech Parameters, in: Introduction to EEG- and Speech-Based Emotion Recognition. Elsevier, pp. 51–79. 10.1016/B978-0-12-804490-2.00003-8

Aftanas, L.I., Golocheikine, S.A., 2001. Human anterior and frontal midline theta and lower alpha reffect emotionally positive state and internalized attention: high-resolution EEG investigation of meditation. Neurosci. Lett. 10.1016/S0304-3940(01)02094-8

Albera, L., Kachenoura, A., Comon, P., Karfoul, A., Wendling, F., Senhadji, L., Merlet, I., 2012. ICA-based EEG denoising: a comparative analysis of fifteen methods. Bull. Pol. Acad. Sci. Tech. Sci. 60, 407–418. 10.2478/v10175-012-0052-3

Ammar, A., Boujelbane, M., Simak, M., Fraile-Fuente, I., Rizz, N., Washif, J., Zmijewski, P., Jahrami, H., Schöllhorn, W., 2024. Unveiling the acute neurophysiological responses to strength training: an exploratory study on novices performing weightlifting bouts with different motor learning models. Biol. Sport. 10.5114/biol-sport.2024.133481

Anders, P., Lehmann, T., Müller, H., Grønvik, K.B., Skjæret-Maroni, N., Baumeister, J., Vereijken, B., 2018. Exergames Inherently Contain Cognitive Elements as Indicated by Cortical Processing. Front. Behav. Neurosci. 12, 102. 10.3389/fnbeh.2018.00102

Baddeley, A.D., 2018. Exploring working memory: selected works of Alan Baddeley, World library of psychologists. Routledge, London.

Baddeley, A.D., Hitch, G., 1974. Working Memory, in: Psychology of Learning and Motivation. Elsevier, pp. 47–89. 10.1016/S0079-7421(08)60452-1

Ball, T., Demandt, E., Mutschler, I., Neitzel, E., Mehring, C., Vogt, K., Aertsen, A., Schulze-Bonhage, A., 2008. Movement related activity in the high gamma range of the human EEG. NeuroImage 41, 302–310. 10.1016/j.neuroimage.2008.02.032

Baumeister, J., Reinecke, K., Cordes, M., Lerch, C., Weiß, M., 2010. Brain activity in goal-directed movements in a real compared to a virtual environment using the Nintendo Wii. Neurosci. Lett. 481, 47–50. 10.1016/j.neulet.2010.06.051

Baumeister, J., Reinecke, K., Liesen, H., Weiss, M., 2008. Cortical activity of skilled performance in a complex sports related motor task. Eur. J. Appl. Physiol. 104, 625–631. 10.1007/s00421-008-0811-x

Becker, L., Büchel, D., Lehmann, T., Kehne, M., Baumeister, J., 2023. Mobile Electroencephalography Reveals Differences in Cortical Processing During Exercises With Lower and Higher Cognitive Demands in Pre-adolescent Children. Pediatr. Exerc. Sci. 35, 214–224. 10.1123/pes.2021-0212

Benedek, M., Schickel, R.J., Jauk, E., Fink, A., Neubauer, A.C., 2014. Alpha power increases in right parietal cortex reflects focused internal attention. Neuropsychologia 56, 393–400. 10.1016/j.neuropsychologia.2014.02.010

Ben-Soussan, T.D., Glicksohn, J., Goldstein, A., Berkovich-Ohana, A., Donchin, O., 2013. Into the Square and out of the Box: The effects of Quadrato Motor Training on Creativity and Alpha Coherence. PLoS ONE 8, e55023. 10.1371/journal.pone.0055023

Benzing, V., Heinks, T., Eggenberger, N., Schmidt, M., 2016. Acute Cognitively Engaging Exergame-Based Physical Activity Enhances Executive Functions in Adolescents. PLOS ONE 11, e0167501. 10.1371/journal.pone.0167501

Boere, K., Lloyd, K., Binsted, G., Krigolson, O.E., 2023. Exercising is good for the brain but exercising outside is potentially better. Sci. Rep. 13, 1140. 10.1038/s41598-022-26093-2

Boutcher, S.H., Landers, D.M., 1988. The Effects of Vigorous Exercise on Anxiety, Heart Rate, and Alpha Activity of Runners and Nonrunners. Psychophysiology 25, 696–702. 10.1111/j.1469-8986.1988.tb01911.x

Büchel, D., Sandbakk, Ø., Baumeister, J., 2021. Exploring intensity-dependent modulations in EEG resting-state network efficiency induced by exercise. Eur. J. Appl. Physiol. 121, 2423–2435. 10.1007/s00421-021-04712-6

Büchel, D., Torvik, P.Ø., Lehmann, T., Sandbakk, Ø., Baumeister, J., 2023. The Mode of Endurance Exercise Influences Changes in EEG Resting-State Graphs among High-Level Cross-Country Skiers. Med. Sci. Sports Exerc. 55, 1003–1013. 10.1249/MSS.0000000000003122

Budde, H., Voelcker-Rehage, C., Pietraßyk-Kendziorra, S., Ribeiro, P., Tidow, G., 2008. Acute coordinative exercise improves attentional performance in adolescents. Neurosci. Lett. 441, 219–223. 10.1016/j.neulet.2008.06.024

Buschman, T.J., Miller, E.K., 2007. Top-Down Versus Bottom-Up Control of Attention in the Prefrontal and Posterior Parietal Cortices. Science 315, 1860–1862. 10.1126/science.1138071

Cavanagh, J.F., Frank, M.J., 2014. Frontal theta as a mechanism for cognitive control. Trends Cogn. Sci. 18, 414–421. 10.1016/j.tics.2014.04.012

Cavanagh, J.F., Shackman, A.J., 2015. Frontal midline theta reflects anxiety and cognitive control: Meta-analytic evidence. J. Physiol.-Paris 109, 3–15. 10.1016/j.jphysparis.2014.04.003

Chang, L.-J., Lin, J.-F., Lin, C.-F., Wu, K.-T., Wang, Y.-M., Kuo, C.-D., 2011. Effect of body position on bilateral EEG alterations and their relationship with autonomic nervous modulation in normal subjects. Neurosci. Lett. 490, 96–100. 10.1016/j.neulet.2010.12.034

Chu, D., Chen, L.-J., Lee, Y.-L., Hung, B.-L., Chou, K.-M., Sun, A.-C., Fang, S.-H., 2018. The correlation of brainwaves of Taekwondo athletes with training vis-à-vis competition performance –an explorative study. Int. J. Perform. Anal. Sport 18, 69–77. 10.1080/24748668.2018.1447205

Corbetta, M., Shulman, G.L., 2002. Control of goal-directed and stimulus-driven attention in the brain. Nat. Rev. Neurosci. 3, 201–215. 10.1038/nrn755

Corsi-Cabrera, M., Ramos, J., Guevara, M.A., Arce, C., Gutierrez, S., 1993. Gender Differencesm in the Eeg During Cognitive Activity. Int. J. Neurosci. 72, 257–264. 10.3109/00207459309024114

Crabbe, J.B., Dishman, R.K., 2004. Brain electrocortical activity during and after exercise: A quantitative synthesis. Psychophysiology 41, 563–574. 10.1111/j.1469-8986.2004.00176.x

Deiber, M.-P., Missonnier, P., Bertrand, O., Gold, G., Fazio-Costa, L., Ibañez, V., Giannakopoulos, P., 2007. Distinction between Perceptual and Attentional Processing in Working Memory Tasks: A Study of Phase-locked and Induced Oscillatory Brain Dynamics. J. Cogn. Neurosci. 19, 158–172. 10.1162/jocn.2007.19.1.158

Doppelmayr, M., Finkenzeller, T., Sauseng, P., 2008. Frontal midline theta in the pre-shot phase of rifle shooting: Differences between experts and novices. Neuropsychologia 46, 1463–1467. 10.1016/j.neuropsychologia.2007.12.026

Engel, A.K., Fries, P., 2010. Beta-band oscillations — signalling the status quo? Curr. Opin. Neurobiol. 20, 156–165. 10.1016/j.conb.2010.02.015

Ermutlu, N., Yücesir, I., Eskikurt, G., Temel, T., İşoğlu-Alkaç, Ü., 2015. Brain electrical activities of dancers and fast ball sports athletes are different. Cogn. Neurodyn. 9, 257–263. 10.1007/s11571-014-9320-2

Fang, Q., Fang, C., Li, L., Song, Y., 2022. Impact of sport training on adaptations in neural functioning and behavioral performance: A scoping review with meta-analysis on EEG research. J. Exerc. Sci. Fit. 20, 206–215. 10.1016/j.jesf.2022.04.001

Fernandes, J., Arida, R.M., Gomez-Pinilla, F., 2017. Physical exercise as an epigenetic modulator of brain plasticity and cognition. Neurosci. Biobehav. Rev. 80, 443–456. 10.1016/J.NEUBIO-REV.2017.06.012

Gebel, A., Lehmann, T., Granacher, U., 2020. Balance task difficulty affects postural sway and cortical activity in healthy adolescents. Exp. Brain Res. 238, 1323–1333. 10.1007/s00221-020-05810-1

Gevins, A., 1997. High-resolution EEG mapping of cortical activation related to working memory: effects of task difficulty, type of processing, and practice. Cereb. Cortex 7, 374–385. 10.1093/cer-cor/7.4.374

Gola, M., Magnuski, M., Szumska, I., Wróbel, A., 2013. EEG beta band activity is related to attention and attentional deficits in the visual performance of elderly subjects. Int. J. Psychophysiol. 89, 334–341. 10.1016/j.ijpsycho.2013.05.007

Gramkow, M.H., Hasselbalch, S.G., Waldemar, G., Frederiksen, K.S., 2020. Resting State EEG in Exercise Intervention Studies: A Systematic Review of Effects and Methods. Front. Hum. Neurosci. 14, 155. 10.3389/fnhum.2020.00155

Guan, A., Wang, S., Huang, A., Qiu, C., Li, Y., Li, X., Wang, J., Wang, Q., Deng, B., 2022. The role of gamma oscillations in central nervous system diseases: Mechanism and treatment. Front. Cell. Neurosci. 16, 962957. 10.3389/fncel.2022.962957

Gutmann, B., Zimmer, P., Hülsdünker, T., Lefebvre, J., Binnebößel, S., Oberste, M., Bloch, W., Strüder, H.K., Mierau, A., 2018. The effects of exercise intensity and post-exercise recovery time on cortical activation as revealed by EEG alpha peak frequency. Neurosci. Lett. 668, 159–163. 10.1016/j.neulet.2018.01.007

Harmony, T., 2013. The functional significance of delta oscillations in cognitive processing. Front. Integr. Neurosci. 7. 10.3389/fnint.2013.00083

Heilmann, F., Weinberg, H., Wollny, R., 2022. The Impact of Practicing Open-vs. Closed-Skill Sports on Executive Functions—A Meta-Analytic and Systematic Review with a Focus on Characteristics of Sports. Brain Sci. 12, 1071. 10.3390/brainsci12081071

Henz, D., John, A., Merz, C., Schöllhorn, W.I., 2018. Post-task Effects on EEG Brain Activity Differ for Various Differential Learning and Contextual Interference Protocols. Front. Hum. Neurosci. 12, 19. 10.3389/fnhum.2018.00019

Henz, D., Schöllhorn, W.I., 2016. Differential Training Facilitates Early Consolidation in Motor Learning. Front. Behav. Neurosci. 10. 10.3389/fnbeh.2016.00199

Herrmann, C.S., Mecklinger, A., 2001. Gamma activity in human EEG is related to highspeed memory comparisons during object selective attention. Vis. Cogn. 8, 593–608. 10.1080/13506280143000142

Herweg, N.A., Solomon, E.A., Kahana, M.J., 2020. Theta Oscillations in Human Memory. Trends Cogn. Sci. 24, 208–227. 10.1016/j.tics.2019.12.006

Hosang, L., Mouchlianitis, E., Guérin, S.M.R., Karageorghis, C.I., 2022. Effects of exercise on electroencephalography-recorded neural oscillations: a systematic review. Int. Rev. Sport Exerc. Psychol. 1–54. 10.1080/1750984X.2022.2103841

Hottenrott, K., Taubert, M., Gronwald, T., 2013. Cortical Brain Activity is Influenced by Cadence in Cyclists. Open Sports Sci. J. 6, 9–14. 10.2174/1875399×01306010009

Hülsdünker, T., Mierau, A., Neeb, C., Kleinöder, H., Strüder, H.K., 2015. Cortical processes associated with continuous balance control as revealed by EEG spectral power. Neurosci. Lett. 592, 1–5. 10.1016/j.neulet.2015.02.049

Hülsdünker, T., Mierau, A., Strüder, H.K., 2016. Higher Balance Task Demands are Associated with an Increase in Individual Alpha Peak Frequency. Front. Hum. Neurosci. 9. 10.3389/fnhum.2015.00695

Jensen, O., Kaiser, J., Lachaux, J.-P., 2007. Human gamma-frequency oscillations associated with attention and memory. Trends Neurosci. 30, 317–324. 10.1016/j.tins.2007.05.001

Jensen, O., Tesche, C.D., 2002. Frontal theta activity in humans increases with memory load in a working memory task. Eur. J. Neurosci. 15, 1395–1399. 10.1046/j.1460-9568.2002.01975.x

John, A.T., Barthel, A., Wind, J., Rizzi, N., Schöllhorn, W.I., 2022. Acute Effects of Various Movement Noise in Differential Learning of Rope Skipping on Brain and Heart Recovery Analyzed by Means of Multiscale Fuzzy Measure Entropy. Front. Behav. Neurosci. 16, 816334. 10.3389/fnbeh.2022.816334

John, A.T., Schöllhorn, W.I., 2018. Acute Effects of Instructed and Self-Created Variable Rope Skipping on EEG Brain Activity and Heart Rate Variability. Front. Behav. Neurosci. 12, 311. 10.3389/fnbeh.2018.00311

John, A.T., Wind, J., Horst, F., Schöllhorn, W.I., 2020. Acute Effects of an Incremental Exercise Test on Psychophysiological Variables and Their Interaction. J. Sports Sci. Med. 19, 596–612.

Jung, J.-Y., Cho, H.-Y., Kang, C.-K., 2020. Brain activity during a working memory task in different postures: an EEG study. Ergonomics 63, 1359–1370. 10.1080/00140139.2020.1784467

Kahya, M., Gouskova, N.A., Lo, O.-Y., Zhou, J., Cappon, D., Finnerty, E., Pascual-Leone, A., Lipsitz, L.A., Hausdorff, J.M., Manor, B., 2022. Brain activity during dual-task standing in older adults. J. NeuroEngineering Rehabil. 19, 123. 10.1186/s12984-022-01095-3

Kao, S.-C., Huang, C.-J., Hung, T.-M., 2013. Frontal Midline Theta is a Specific Indicator of Optimal Attentional Engagement During Skilled Putting Performance. 10.1123/jsep.35.5.470

Kirschbaum, C. (Ed.), 2008. Biopsychologie von A bis Z: inklusive Online-Version, Springer-Lehrbuch Bachelor, Master. Springer-Medizin-Verl, Heidelberg.

Klimesch, W., 1999. EEG alpha and theta oscillations reflect cognitive and memory performance: a review and analysis. Brain Res. Rev. 29, 169–195. 10.1016/S0165-0173(98)00056-3

Klimesch, W., 1997. EEG-alpha rhythms and memory processes. Int. J. Psychophysiol. 26, 319–340. 10.1016/S0167-8760(97)00773-3

Klimesch, W., Sauseng, P., Hanslmayr, S., 2007. EEG alpha oscillations: The inhibition–timing hypothesis. Brain Res. Rev. 53, 63–88. 10.1016/j.brainresrev.2006.06.003

Kubitz, K.A., Landers, D.M., 1993. The Effects of Aerobic Training on Cardiovascular Responses to Mental Stress: An Examination of Underlying Mechanisms. J. Sport Exerc. Psychol. 15, 326–337. 10.1123/jsep.15.3.326

Lalo, E., Gilbertson, T., Doyle, L., Lazzaro, V.D., Cioni, B., Brown, P., 2007. Phasic increases in cortical beta activity are associated with alterations in sensory processing in the human. Exp. Brain Res. 177, 137–145. 10.1007/s00221-006-0655-8

Lardon, M.T., Polich, J., 1996. EEG changes from long-term physical exercise. Biol. Psychol. 44, 19–30. 10.1016/S0301-0511(96)05198-8

Lau, T.M., Gwin, J.T., Ferris, D.P., 2012. How Many Electrodes Are Really Needed for EEG-Based Mobile Brain Imaging? J. Behav. Brain Sci. 02, 387–393. 10.4236/jbbs.2012.23044

Malik, A.S., Amin, H.U., 2017. Designing EEG experiments for studying the brain: design code and example datasets. Academic Press, an imprint of Elsevier, London San Diego.

Mari-Acevedo, J., Yelvington, K., Tatum, W.O., 2019. Normal EEG variants, in: Handbook of Clinical Neurology. Elsevier, pp. 143–160. 10.1016/B978-0-444-64032-1.00009-6

Mehta, R.K., Parasuraman, R., 2013. Neuroergonomics: a review of applications to physical and cognitive work. Front. Hum. Neurosci. 7. 10.3389/fnhum.2013.00889

Miraglia, F., Tomino, C., Vecchio, F., Alù, F., Orticoni, A., Judica, E., Cotelli, M., Rossini, P.M., 2021. Assessing the dependence of the number of EEG channels in the brain networks’ modulations. Brain Res. Bull. 167, 33–36. 10.1016/j.brainresbull.2020.11.014

Misselhorn, J., Friese, U., Engel, A.K., 2019. Frontal and parietal alpha oscillations reflect attentional modulation of cross-modal matching. Sci. Rep. 9, 5030. 10.1038/s41598-019-41636-w

Müller, H., Baumeister, J., Bardal, E.M., Vereijken, B., Skjæret-Maroni, N., 2023. Exergaming in older adults: the effects of game characteristics on brain activity and physical activity. Front. Aging Neurosci. 15, 1143859. 10.3389/fnagi.2023.1143859

National Collaborating Centre for Methods and Tools [WWW Document], 2008.. Qual. Assess. Tool Quant. Stud. URL https://www.nccmt.ca/knowledge-repositories/search/14 (accessed 5.15.24).

Noudoost, B., Chang, M.H., Steinmetz, N.A., Moore, T., 2010. Top-down control of visual attention. Curr. Opin. Neurobiol. 20, 183–190. 10.1016/j.conb.2010.02.003

Palva, S., Palva, J.M., 2007. New vistas for α-frequency band oscillations. Trends Neurosci. 30, 150–158. 10.1016/j.tins.2007.02.001

Parr, J.V.V., Gallicchio, G., Wood, G., 2021. EEG correlates of verbal and conscious processing of motor control in sport and human movement: a systematic review. Int. Rev. Sport Exerc. Psychol. 16, 396–427. 10.1080/1750984X.2021.1878548

Pesce, C., 2012. Shifting the Focus From Quantitative to Qualitative Exercise Characteristics in Exercise and Cognition Research. J. Sport Exerc. Psychol. 34, 766–786. 10.1123/jsep.34.6.766

Piantoni, G., Kline, K.A., Eagleman, D.M., 2010. Beta oscillations correlate with the probability of perceiving rivalrous visual stimuli. J. Vis. 10, 18–18. 10.1167/10.13.18

Rutkove, S.B., 2007. Introduction to Volume Conduction, in: Blum, A.S., Rutkove, S.B. (Eds.), The Clinical Neurophysiology Primer. Humana Press, Totowa, NJ, pp. 43–53. 10.1007/978-1-59745-271-7_4

Rydzik, Ł., Obmiński, Z., Wąsacz, W., Kopańska, M., Kubacki, R., Bagińska, M., Tota, Ł., Ambroży, T., Witkowski, K., Pałka, T., 2024. The effect of physical exercise during competitions and in simulated conditions on hormonal-neurophysiological relationships in kickboxers. Biol. Sport 41, 61–68. 10.5114/biolsport.2024.133662

Sauseng, P., Hoppe, J., Klimesch, W., Gerloff, C., Hummel, F.C., 2007. Dissociation of sustained attention from central executive functions: local activity and interregional connectivity in the theta range. Eur. J. Neurosci. 25, 587–593. 10.1111/j.1460-9568.2006.05286.x

Sibley, B.A., Etnier, J.L., 2003. The relationship between physical activity and cognition in children: A meta-analysis. Pediatr. Exerc. Sci. 15, 243–256. 10.1123/PES.15.3.243

Sipp, A.R., Gwin, J.T., Makeig, S., Ferris, D.P., 2013. Loss of balance during balance beam walking elicits a multifocal theta band electrocortical response. J. Neurophysiol. 110, 2050–2060. 10.1152/jn.00744.2012

Teplan, M., 2002. FUNDAMENTALS OF EEG MEASUREMENT. Meas. Sci. Rev. 2, 1–11.

The EndNote Team, 2013. EndNote.

Thompson, L., Khuc, J., Saccani, M.S., Zokaei, N., Cappelletti, M., 2021. Gamma oscillations modulate working memory recall precision. Exp. Brain Res. 239, 2711–2724. 10.1007/s00221-021-06051-6

Tomporowski, P.D., 2003. Effects of acute bouts of exercise on cognition. Acta Psychol. (Amst.) 112, 297–324. 10.1016/S0001-6918(02)00134-8

Tricco, A.C., Lillie, E., Zarin, W., O’Brien, K.K., Colquhoun, H., Levac, D., Moher, D., Peters, M.D.J., Horsley, T., Weeks, L., Hempel, S., Akl, E.A., Chang, C., McGowan, J., Stewart, L., Hartling, L., Aldcroft, A., Wilson, M.G., Garritty, C., Lewin, S., Godfrey, C.M., Macdonald, M.T., Langlois, E.V., Soares-Weiser, K., Moriarty, J., Clifford, T., Tunçalp, Ö., Straus, S.E., 2018. PRISMA Extension for Scoping Reviews (PRISMA-ScR): Checklist and Explanation. Ann. Intern. Med. 169, 467–473. 10.7326/M18-0850

Tuladhar, A.M., Huurne, N. ter, Schoffelen, J., Maris, E., Oostenveld, R., Jensen, O., 2007. Parieto-occipital sources account for the increase in alpha activity with working memory load. Hum. Brain Mapp. 28, 785–792. 10.1002/hbm.20306

Ulloa, J.L., 2022. The Control of Movements via Motor Gamma Oscillations. Front. Hum. Neurosci. 15, 787157. 10.3389/fnhum.2021.787157

van Schouwenburg, M.R., Zanto, T.P., Gazzaley, A., 2017. Spatial Attention and the Effects of Frontoparietal Alpha Band Stimulation. Front. Hum. Neurosci. 10. 10.3389/fnhum.2016.00658

Varela, F., Lachaux, J.-P., Rodriguez, E., Martinerie, J., 2001. The brainweb: Phase synchronization and large-scale integration. Nat. Rev. Neurosci. 2, 229–239. 10.1038/35067550

Visser, A., Büchel, D., Lehmann, T., Baumeister, J., 2022. Continuous table tennis is associated with processing in frontal brain areas: an EEG approach. Exp. Brain Res. 240, 1899–1909. 10.1007/s00221-022-06366-y

Wind, J., Horst, F., Rizzi, N., John, A., Schöllhorn, W.I., 2020. Electrical Brain Activity and Its Functional Connectivity in the Physical Execution of Modern Jazz Dance. Front. Psychol. 11, 586076. 10.3389/fpsyg.2020.586076

Wittenberg, E., Thompson, J., Nam, C.S., Franz, J.R., 2017. Neuroimaging of Human Balance Control: A Systematic Review. Front. Hum. Neurosci. 11. 10.3389/fnhum.2017.00170

Wollseiffen, P., Ghadiri, A., Scholz, A., Strüder, H.K., Herpers, R., Peters, T., Schneider, S., 2016. Short Bouts of Intensive Exercise During the Workday Have a Positive Effect on Neuro-cognitive Performance. Stress Health. 10.1002/smi.2654

Wu, Q., Jiang, H., Shao, C., Zhang, Y., Zhou, W., Cao, Y., Song, J., Shi, B., Chi, A., Wang, C., 2023. Characteristics of changes in the functional status of the brain before and after 1,000 m all-out paddling for different levels of dragon boat athletes. Front. Psychol. 14, 1109949. 10.3389/fpsyg.2023.1109949

Zschocke, S., Kursawe, H., Kursawe, H.K. (Eds.), 2012. Klinische Elektroenzephalographie: mit DVD: [EEG-Beispiele zum Auswerten], 3., aktualisierte und erw. Aufl. ed. Springer, Berlin Heidelberg.

